# Glycolytic and serine biosynthetic pathways with a novel mitochondrial location contribute to fitness in the oomycete *Phytophthora*

**DOI:** 10.64898/2026.07.27.741032

**Authors:** Carl S. Mendoza, Howard S. Judelson

## Abstract

The eukaryotic microbes known as oomycetes express both cytosolic and mitochondrial sets of enzymes for the last six steps in glycolysis, unlike canonical eukaryotes where glycolysis is exclusively cytosolic. Linked to glycolysis through 3-phosphoglycerate is another pathway with an atypical mitochondrial location in oomycetes, phosphorylated serine biosynthesis (PSB), which is cytosolic in most other eukaryotes. Previous studies confirmed the mitochondrial location of these enzymes, which are encoded by nuclear genes. However, information was lacking about their contributions to metabolism and fitness. We addressed this using *Phytophthora infestans,* a pathogen of potato and tomato. Single-gene knockouts of the mitochondrial forms of phosphoglycerate kinase and enolase had little effect on growth or pathogenicity, but these traits were severely impaired in strains deleted for both genes. Metabolomic analysis of the double knockouts revealed changes in levels of glycolytic and TCA cycle intermediates, adenylate pools, vitamins, and other compounds important to cellular function. Blocking the PSB pathway by knocking out phosphoserine aminotransferase also compromised growth but caused fewer metabolic changes, and the results suggested that the main role of the pathway is to generate 3-phosphoglycerate for glycolysis and not for making serine. Most of these enzymes appear to have been acquired by lateral gene transfer into the stramenopile lineage, which besides oomycetes include diatoms and brown algae. We conclude that at least in oomycetes, metabolism has adapted such that these enzymes are now required for fitness.

**IMPORTANCE:** The non-sexual movement of genetic material between species (horizontal gene transfer, HGT) has driven evolution in both prokaryotes and eukaryotes. One impact of HGT on the eukaryotic microbes known as oomycetes has been the acquisition of a glycolytic pathway in mitochondria alongside the standard version that resides in the cytosol. This is an uncommon outcome of HGT involving metabolic genes, which usually results in replacement of the ancestral pathway. We show that the mitochondrial glycolytic pathway, and another novel pathway that shares a metabolic intermediate with glycolysis in that organelle, make important contributions to metabolism and are required for fitness. This study helps illustrate how oomycete metabolism has adapted to the acquisition of enzymes with different evolutionary origins. Also, since oomycetes cause diseases on many important crop plants and animals, these novel features of metabolism may provide targets for their chemical control.

## INTRODUCTION

Membrane-bound organelles are a defining feature of eukaryotic cells (1). Many organelles are thought to have evolved via endosymbiosis, in which free-living microbes were engulfed by an ancestral host cell (2, 3). One consequence is the compartmentalization of metabolism. By sequestering metabolic pathways within an organelle cells can optimize reaction conditions, increase local substrate and enzyme concentrations, establish gradients to drive reactions, and prevent futile cycling (4, 5). Mitochondria exemplify compartmentalization and serve as a central hub for ATP production and the metabolism of carbohydrates, lipids, and amino acids through the tricarboxylic acid (TCA) cycle and other pathways (6).

In contrast to the TCA cycle which resides exclusively in mitochondria, glycolysis is cytosolic in most eukaryotes (7). Glycolysis begins with a so-called preparatory phase involving four enzymes that convert glucose to triose phosphates at the expense of ATP. This is followed by a pay-off phase of six enzymes that generate pyruvate with a net gain of ATP (Fig. 1A). Pyruvate may then be oxidized and form acetyl-CoA to enter the TCA cycle. However, pyruvate must first be transported into the mitochondrial matrix, which is considered to be a rate-limiting step in energy production (8). Glycolysis is also linked to other pathways such as the phosphorylated serine biosynthesis (PSB) pathway (Fig. 1A). Depending on cellular needs, this either uses the pay-off phase intermediate 3-phosphoglycerate to make serine (9) or to feed carbon from serine into glycolysis or gluconeogenesis (10).

**Fig. 1.**
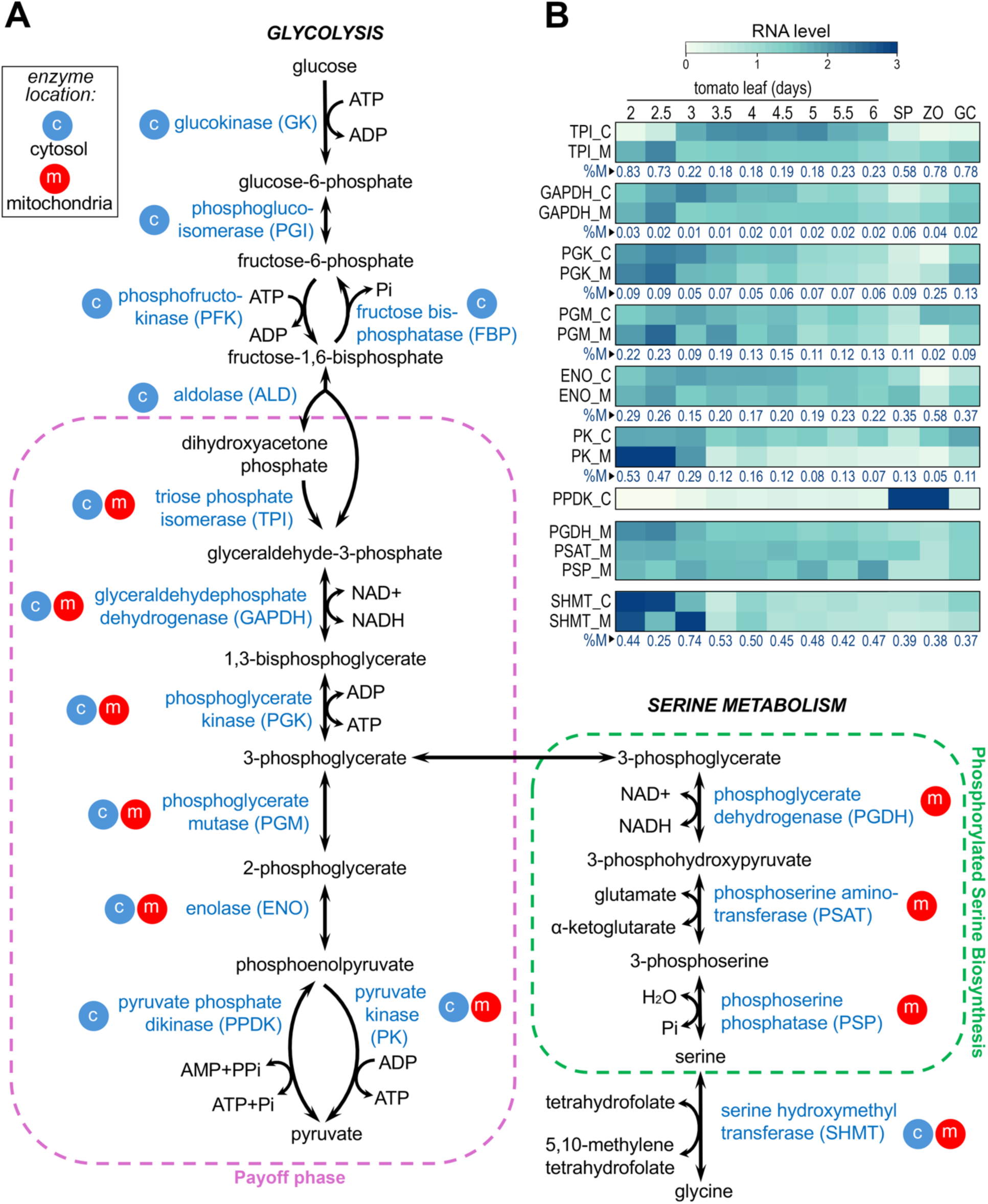
Glycolysis and serine biosynthesis pathways in *P. infestans.* **A,** Diagrams showing whether the enzymes localize to the cytosol or mitochondria, as indicated by the c and m emblems, respectively. The ATP-generating pay-off phase of glycolysis is marked by the purple dashed border and the phosphorylated serine biosynthesis (PSB) pathway by the green dashed line. Also shown is the SHMT pathway for making serine. **B,** mRNA levels of the genes for the enzymes from the pay-off phase and the serine pathways during growth in tomato leaves and in purified sporangia (SP), zoospores (Z), and germinated zoospore cysts (GC). Values are per-gene normalized such that the mean of each row equals 1.0. Enzyme names are appended with C or M to denote the cytosolic or mitochondrial activities, respectively. Marked beneath each subpanel percentage are t he FPKM values from the mitochondrial genes compared to the total (%M). If multiple genes encoded the same enzyme, their FPKM values were combined. FPKM values of each gene are shown in Table S1.

While glycolysis is normally cytosolic there are a few exceptions. In kinetoplastid protozoa such as *Trypanosoma brucei,* the first seven glycolytic enzymes are located inside the peroxisome-like organelle called the glycosome, while the last three are cytosolic (11). In the apicomplexan *Toxoplasma gondii,* four of the six canonical pay-off phase enzymes have isoforms that reside in the apicoplast, a plastid-like organelle (12). In plants, up to nine of the glycolytic enzymes occur in the chloroplast, in addition to their cytosolic location (13). After the ancient engulfment of a cyanobacterium to form the chloroplast, such genes were transferred to the host nucleus (14).

In prior work with the oomycete *Phytophthora infestans,* a member of the stramenopile group and an important pathogen of potato and tomato, we identified mitochondrial targeting peptides in each of the six glycolytic pay-off phase enzymes. All are encoded by nuclear genes. We then used fluorescent tags to confirm that the proteins reside in mitochondria (15). Thus, this filamentous eukaryotic microbe has a complete set of glycolytic enzymes that reside in the cytosol plus a second set that establishes the pay-off phase in the mitochondria. We also observed that the three enzymes of the PSB pathway (phosphoserine aminotransferase, phosphoglycerate dehydrogenase, and phosphoserine phosphatase) are also mitochondrial in *P. infestans,* while being cytosolic in canonical eukaryotes. Genome mining indicated that these novel features are shared by other stramenopiles including diatoms, brown algae, and other oomycetes, and likely appeared through endosymbiotic or horizontal gene transfer (E/HGT). A subsequent study in diatoms also confirmed that their mitochondria also contain the pay-off phase enzymes (16).

What was lacking from our prior study was evidence that the mitochondrial glycolytic and serine pathways contribute significantly to metabolism, as opposed to being functionally redundant. This was an open question since proteomic data from *P. infestans* suggested that the cytosolic pay-off phase enzymes are more abundant than their mitochondrial counterparts (15). In this study, we fill this knowledge gap by demonstrating that several of the mitochondrial enzymes are sometimes expressed at higher levels than the cytosolic enzymes, and by testing knockouts of genes in the pay-off and PSB pathways. While biallelic knockouts (oomycetes are diploid) of the mitochondrial forms of phosphoglycerate kinase or enolase individually had limited effects on growth, spore production, or virulence, these traits were severely degraded when genes for both enzymes were disrupted in the same strain. Knockouts of the PSB pathway also reduced fitness. Metabolomic analysis identified disruptions in the normal balance of key intermediates in glycolysis, the TCA cycle, serine, and other compounds in the mutants. These findings demonstrate the importance of the mitochondrial enzymes and the integration of mitochondrial glycolysis with the serine pathway, revealing a unique metabolic adaptation in stramenopiles.

## RESULTS

### Mitochondrial and cytosolic glycolytic enzymes exhibit distinct expression patterns

We previously performed a limited analysis of glycolytic gene expression using hyphae from rye grain broth, minimal media, and potato tuber slice cultures (15). To understand expression under less artificial conditions, here we examined a time course of infection on tomato leaves as well as sporangia, zoospores, and germinated zoospore cysts, and extended the analysis to the PSB genes. While our prior study concluded that the genes encoding the mitochondrial glycolytic enzymes were transcribed at much lower levels than their cytosolic paralogs, the new data revealed a more complicated pattern.

During the first two days of plant colonization, several mitochondrial glycolytic genes had higher mRNA levels than their cytosolic counterparts (Fig. 1B). This stage corresponds to the biotrophic period when *P. infestans* grows mostly in the apoplast and forms haustoria. For example, FKPM values for the mitochondrial triose phosphate isomerase (TPI) and pyruvate kinase (PK) genes represented 83% and 53% of the total for those activities, respectively. In contrast, the mitochondria-targeted phosphoglycerate kinase (PGK) and enolase (ENO) genes accounted for only 22 and 29% of the FPKM for those activities, respectively. These patterns changed as infection proceeded. By day five, for example, transcripts for the mitochondrial TPI and PK proteins which earlier were in the majority declined to 18 and 8% of the total, respectively. This timepoint represents the necrotrophic stage of infection when many host cells would be compromised and releasing their metabolites.

In the spore stages, the glycolytic genes also exhibited diverse patterns. Transcripts of both cytosolic and mitochondrial forms were generally lower in sporangia and zoospores compared to the early stages of infection but increased in germinating zoospore cysts. In most cases, mRNA levels were higher for the cytosolic than the mitochondrial proteins. The exception was TPI, for which the FPKM value of its mitochondrial form represented 58, 78, and 78% of the total for TPI in sporangia, zoospores, and germinated cysts, respectively. Mitochondrial ENO mRNA also was slightly more abundant than that of cytosolic ENO in zoospores.

The genes for serine biosynthesis also showed dynamic patterns of expression. For the PSB pathway, these were phosphoserine phosphatase (PSP) and a fusion of phosphoserine aminotransferase (PSAT) and phosphoglycerate dehydrogenase (PGDH); unlike most species where PGDH and PSAT are separate proteins, they are fused in oomycetes and brown algae (15). Transcript levels of these genes were relatively constant during tomato leaf infection, declined in sporangia and zoospores, and then increases in germinating cysts. While the PSB genes only have mitochondrial forms in *P. infestans,* serine hydroxymethyltransferase (SHMT), which makes serine from glycine, has both mitochondrial and cytosolic forms. mRNA levels of both SHMT genes declined as the infection proceeded, and these diminished levels persisted in the spore stages. The proportion of transcripts for mitochondrial and cytosolic SHMT were generally balanced.

### The mitochondrial glycolytic pathway contributes to fitness

Gene knockouts generated by CRISPR-Cas12a were used to investigate the role of the mitochondrial glycolytic pay-off phase. Initially, we targeted separately the mitochondrial forms of phosphoglycerate kinase, phosphoglycerate mutase, and enolase, referred to hereafter as mtPGK, mtPGM, and mtENO, respectively. We obtained strains with biallelic knockouts for mtPGK (PITG_00132) at a frequency of 3 out of 100 transformants, and for mtENO (PITG_14195) in 3 out of 130 transformants. These were initially identified by using polymerase chain reaction to detect deletions (Fig. 2A) and then confirmed for their inability to express a functional protein by Sanger sequencing. We also attempted to identify knockouts of mtPGM, but none were observed among 80 transformants.

**Fig. 2.**
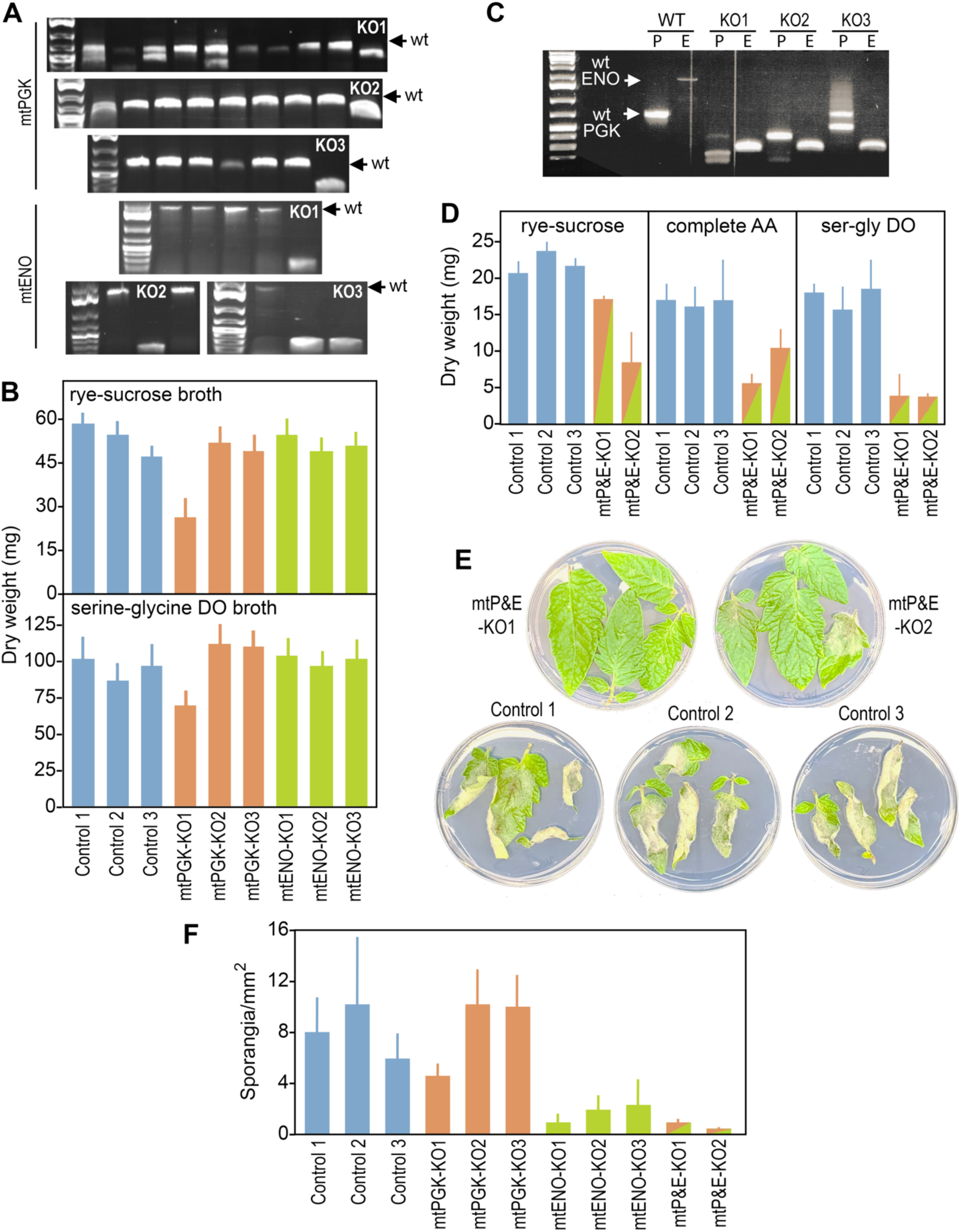
Strains with knockouts of glycolytic enzymes. **A,** Polymerase chain reaction (PCR) screening of primary transformants treated with CRISPR plasmids targeting mtPGK or mtENO. Strains yielding bands with altered mobility compared to wild-type (wt) and selected for later analyses are marked by KO1, KO2, and KO3. **B,** Dry weight growth assays on rye-sucrose (7 days after inoculation) or synthetic amino acid media lacking serine and glycine (DO broth, 20 days). Shown are the three mtPGK knockouts (mtPGK-KO; orange bars), three mtENO knockouts (mtENO-KO; green), and three controls (blue). The latter were the progenitor strain 1306 (Control 1) and two unedited transformants (control 2, control 3). **C,** PCR assays of strains containing knockouts of both mtPGK and mtENO genes (mtP&E-KO), obtained by cotransformation with the two CRISPR plasmids. Shown are three double-knockout strains selected from a prior screen of primary transformants. **D,** Dry weight growth assays after 7, 10, and 10 days on rye-sucrose, synthetic media containing all 20 amino acids, or amino acid media lacking serine and glycine (DO broth), respectively. Only two knockout strains (green/orange bars) are shown since the third died before the experiment could be completed. **E,** Virulence assay of the double-knockout strains on tomato leaves, six days after inoculation. The successfully colonized leaves, which are only seen in the controls, are fuzzy and slightly curled. **F,** Sporangia production of control, single-knockout and double-knockout strains on rye-sucrose agar.

Neither the mtPGK or mtENO knockout strains displayed measurable changes in growth on rich or defined media or virulence on leaves (Fig. 2B). However, the mtENO knockouts produced fewer sporangia than the unedited controls (*P*=0.006), although each produced normal zoospores. The other traits showed some variation between the three knockout strains (*e.g.,* mtPGK-KO1, KO2, and KO3) which we attribute to off-target editing or mutagenic effects of transformation,

Given the existence of the cytosolic pay-off phase, the bidirectionality of glycolysis, and the connection between PSB and the mitochondrial pay-off phase we hypothesized that *P. infestans* might have been able to adapt to the loss of just mtPGK or mtENO. For example, the loss of mtPGK might have been offset by feeding 3-phosphoglycerate from the PSB pathway into the pay-off phase to continue the flow of carbohydrate towards pyruvate. Similarly, the PSB pathway may have compensated for a mtENO defect if the mitochondrial pay-off phase was contributing to gluconeogenesis. To address this, we generated strains in which genes upstream and downstream of 3-phosphoglycerate in glycolysis were both mutated, by expressing guide RNAs targeting both genes in a single plasmid. After screening 500 transformants, several bearing biallelic deletions within both genes were identified. Two were homokaryons and are labelled KO1 and KO2 in Fig. 2C. Others, such as the one labelled KO3 in Fig. 2C, contained biallelic edits in both genes but retained wild-type bands, suggesting that these were heterokaryons with some unedited nuclei. Attempts to obtain homokaryotic edited derivatives by passage through the single-nuclei zoospore stage were unsuccessful, suggesting that the double knockout reduces fitness. This is consistent with our observation that several of the heterokaryotic strains lost their edited alleles upon subculture.

Phenotypic analysis of the double-knockout strains (labeled as mtP&E-KO in Fig. 2D-F) confirmed their reduced fitness. Although the mutants did not exhibit a consistently reduced rate of radial growth, their hyphal mats were visibly less dense. In dry weight assays, hyphal biomass was reduced significantly in the mutants compared to the controls by 18% on rye media (*P*=0.02), 35% on a synthetic media containing all 20 amino acids (*P*=0.003), and 70% on the synthetic media lacking serine and glycine (*P*=0.0001; Fig. 2D). The latter medium was tested since serine made by the PSB pathway or serine hydroxymethyltransferase might partially rescue the metabolic defect. The double knockouts failed to colonize tomato leaves (Fig. 2E) and produced fewer sporangia than controls (*P*=0.004; Fig. 2F), although the sporangia that were produced released normal zoospores. Overall, these data suggest that while *P. infestans* can compensate for single knockouts of mtPGK or mtENO, it cannot compensate for the loss of both genes and the subsequent block to 3-phosphoglycerate formation in mitochondria.

Since the double knockouts might reduce the production of pyruvate in mitochondria, we tested whether their growth defect would be rescued by adding 10 mM pyruvate to the amino acid-based synthetic media. This did not rescue the defects, which is not surprising since the pyruvate would need to be transported from the cytosol to the mitochondria, which is a rate-limiting step in providing carbon to the TCA cycle (8). This highlights a potential benefit of the mitochondrial pay-off phase, as producing pyruvate in mitochondria would bypass that hurdle.

### mtPGK knockouts do not strongly reduce total PGK activity

Whether the mtPGK knockout reduced total PGK catalytic activity was assessed using an enzyme-linked assay with extracts of hyphae from rye broth cultures. There was not a significant difference between the three controls and three knockout strains (Fig. 3). This may indicate that *P. infestans* compensates for the loss of mtPGK by upregulating the cytosolic enzyme. Alternatively, the change may have been too small to detect above normal biological noise. We previously reported that mtPGK protein comprised only about 9% of total PGK protein in hyphae, although activity of the two PGK proteins may not be proportional to their concentrations (15).

**Fig. 3.**
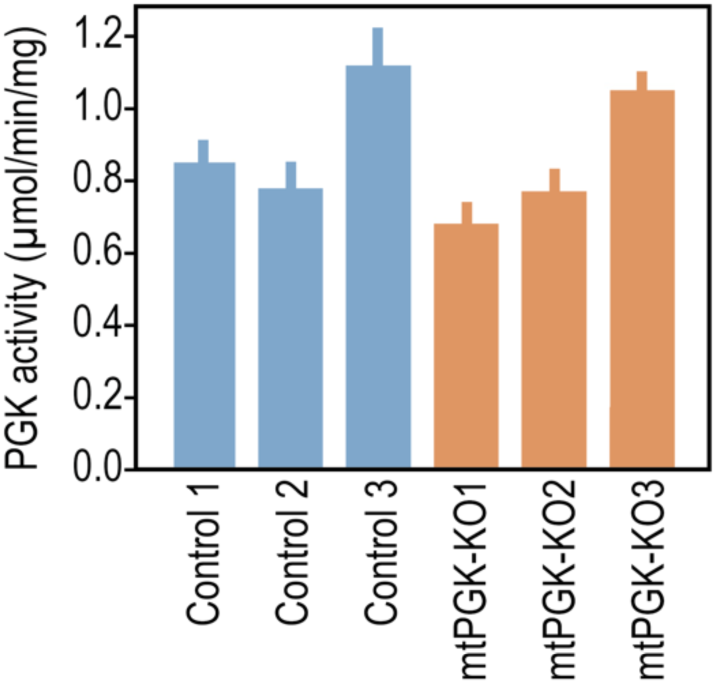
Specific activity of phosphoglycerate kinase in the mtPGK knockout mutants. The controls and knockouts are the same as in Fig. 2.

### Blocking the PSB pathway impairs growth

We extended our analysis to this *de novo* serine biosynthesis pathway by knocking out the gene encoding the PGDH-PSAT fusion, which links the pathway to glycolysis through 3-phosphoglycerate. The gene, PITG_00133, is referred to hereafter as PSAT. Three biallelic knockouts were obtained from 60 transformants (Fig. 4A). Based on dry weight assays, growth of the knockouts was reduced significantly on both rye-sucrose and synthetic media by about 50% (*P=*10^-5^ and *P*=0.003, respectively; Fig. 4B). Like the mtPGK and mtENO double mutants, radial growth of the PSAT knockouts was not consistently diminished relative to controls but their hyphal mats were sparser.

**Fig. 4.**
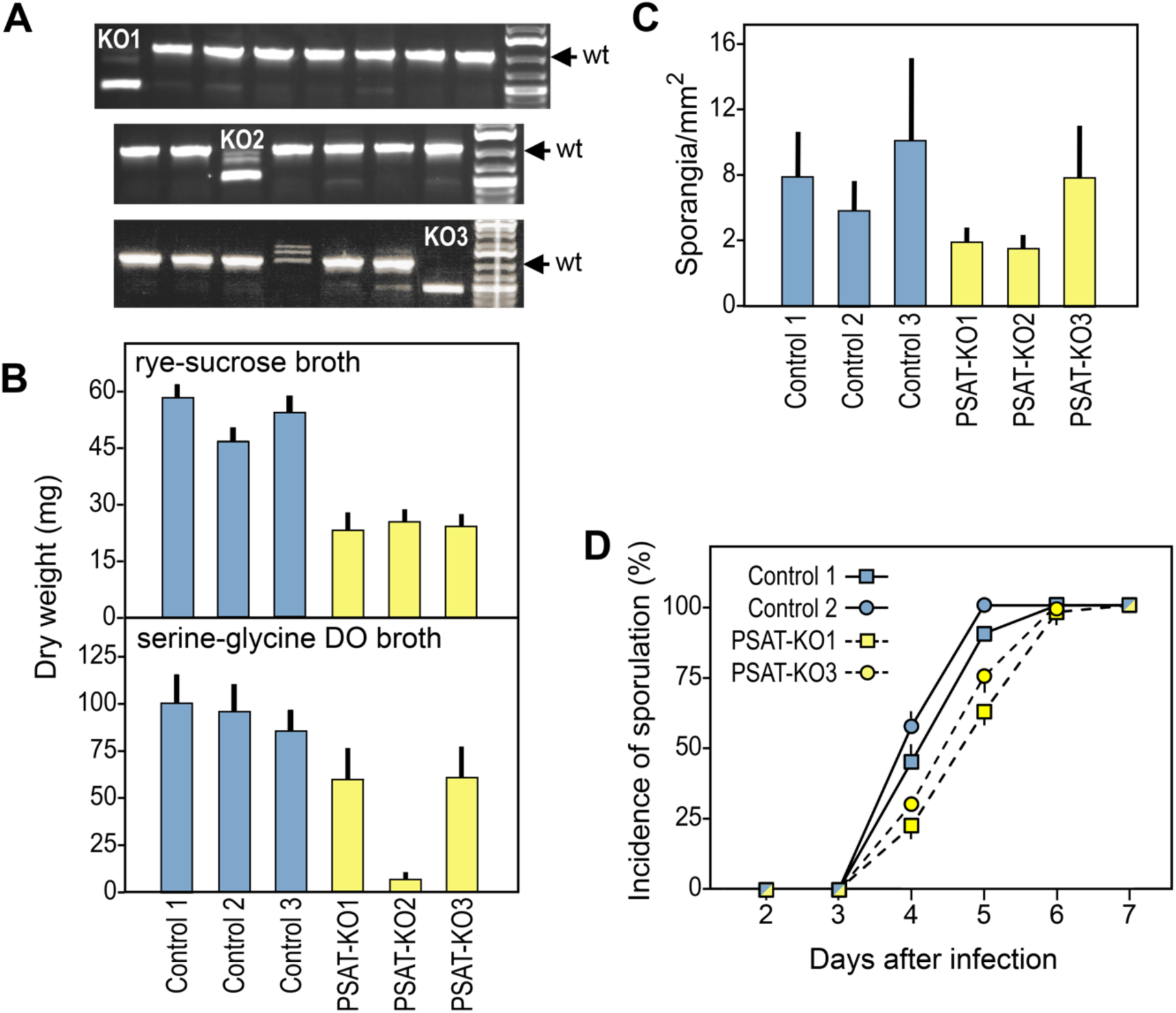
Strains with knockouts of PSAT. **A,** Polymerase chain reaction (PCR) screening of primary transformants treated with CRISPR. Strains yielding bands with altered mobility compared to wild-type (wt) and selected for later analyses are marked KO1, KO2, and KO3. **B,** Dry weight growth assays after 7 days on rye-sucrose broth or 10 days on synthetic amino acid broth lacking serine and glycine (DO broth). Shown are the three mtPSAT knockouts (mtPSAT-KO; yellow bars) and three controls, which are strain 1306 and two unedited transformants (blue bars). **C,** Sporangia production of control and knockout strains on rye-sucrose agar. **D,** Progression of disease on tomato leaflets. Data are the average of two independent experiments, each with 20 leaflets per strain. While sporulation occurred more slowly with the knockouts, there was not a significant difference in the number of sporangia after day 6 (434±56 per mg of infected tissue for the knockouts versus 406±126 for the controls).

Sporulation was not significantly impaired (Fig. 4C), and the sporangia released normal zoospores. However, the PSAT knockouts grew slightly slower on leaves than controls (Fig. 4D). While the knockouts completed the disease cycle, their latent periods (time to sporulation) averaged 5.03±0.76 days compared to 4.54±0.59 for the unedited controls (Fig. 4D). Area Under the Disease Progress Curve calculations (17) indicated that the difference was significant (*P*=0.004) with 19% slower progression of the disease. This was less of a reduction in growth than seen in artificial media, possibly since a leaf provides *P. infestans* with a more suitably balanced set of nutrients.

### mtPGK and mtENO contribute substantially to metabolite pools

To get a clearer picture of the mitochondrial pathways’ roles in metabolite production, we used liquid chromatography-mass spectrometry (LC-MS) to compare central carbon metabolites in the control and knockout strains, using hyphae grown in the amino acid synthetic media lacking serine and glycine. The results for the single knockout mtPGK, mtENO, the double knockout mtP&E (mtPGK & mtENO) strains are discussed here while the PSAT knockouts is addressed in the next section. Selected data are shown in Fig. 5 and the results for all measured compounds are in Table S2.

**Fig. 5.**
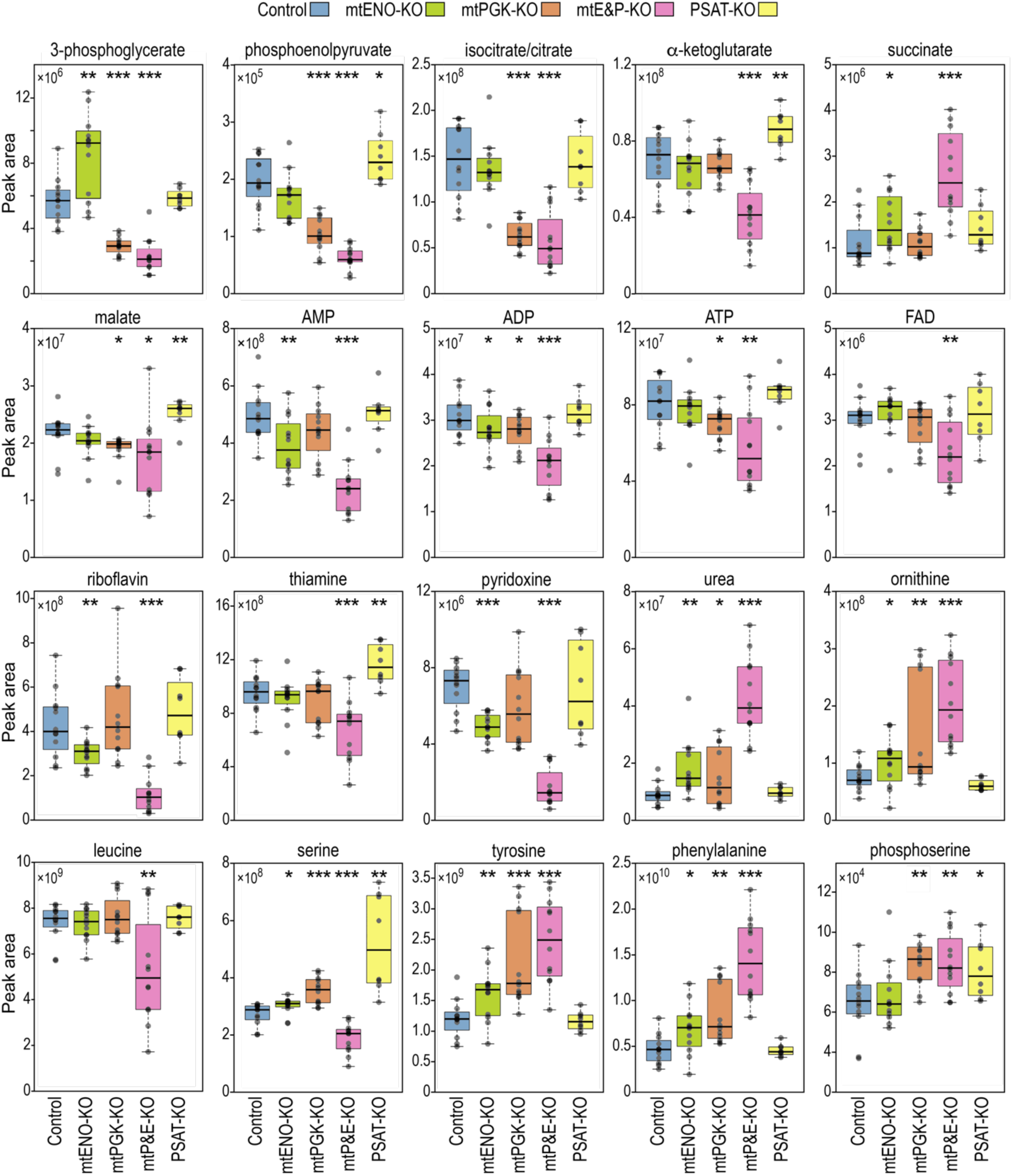
Relative levels of selected metabolites in the single and double-dropout strains, based on targeted metabolomic analysis. Compounds are listed (left to right, top to bottom) in the order mentioned in Results. Other compounds from the analysis are shown in Table S2. Asterisks denote differences with the controls at *P*<0.05 (*), *P*<0.01 (**), and *P*<0.001 (***).

Major differences were observed between the controls and mtPGK knockouts or mtP&E double knockouts, indicating that the mitochondrial pathway contributes substantially to metabolism (Fig. 5). For example, the mtPGK knockouts caused 48 and 47% average declines in the downstream metabolites 3-phosphoglycerate and phosphoenolpyruvate, respectively, while 59% and 72% reductions occurred in the double knockouts. The magnitude of these changes might seem surprising in light of our earlier finding that mtPGK comprised only 9% of total PGK protein (15). However, the turnover number (*K_cat_*) of mtPGK may be higher than that of cytosolic PGK considering that the two proteins are distinct, with only 47% sequence identity (based on a corrected gene model for the cytosolic enzyme, PITG_09402). Moreover, the isoelectric point of the mitochondrial form is higher (6.63 versus 5.68), which may make it better-suited to the more alkaline environment of the mitochondrial matrix (18). mtPGK may also be more active if its substrate concentration is higher within mitochondria, which is a proposed benefit of metabolic compartmentalization (1).

Effects of the knockouts carried through to the TCA cycle, especially for the double knockouts. For example, the mtP&E strains averaged 61% lower levels of the early TCA cycle intermediates isocitrate (indistinguishable from citrate in the analysis) and α-ketoglutarate, likely resulting from reduced delivery of pyruvate. An exception was the mid-cycle intermediate succinate, which rose about two-fold in the mtP&E strains compared to controls. This could reflect a strategy to counter the decline in early intermediates by catabolizing amino acids, as many feed into the cycle through succinyl-CoA (19). Another explanation for the rise in succinate in the knockouts is a bottleneck caused by reduced FAD (discussed below), which is needed to convert succinate to fumarate. Our LC-MS pipeline did not detect fumarate, but malate levels fell by 19%.

The mtENO knockout alone also conferred changes consistent with a functional mitochondrial pay-off phase. This caused a near-doubling of the enolase substrate, 3-phosphoglycerate, and a decline in its phosphoenolpyruvate product (Fig. 5). This is logical if the pay-off phase is operating in the energy-generating direction and not for gluconeogenesis. The drop in phosphoenolpyruvate was not as extreme as the increase in 3-phosphoglycerate, possibly due to generation of the former by pyruvate phosphate dikinase (PPDK). Although our transcription data suggests that the PPDK genes are transcribed mostly in the spore stages (Fig. 1B), a prior study indicated that there is substantial activity in hyphae (20).

Changes were also observed beyond the glycolytic and TCA pathways, especially with the mtP&E knockouts. Many were consistent with mitochondrial dysfunction, such as a drop in the total adenylate pool as 51, 34, and 29% declines were observed in AMP, ADP, and ATP, respectively. Also falling in concentration were B vitamins such as pyridoxine, riboflavin, and thiamine (Fig. 5) and niacin (Table S2). The largest change was for riboflavin, which fell 10-fold in the double knockouts. This is the precursor for FAD, which declined in the double knockouts by 25%. FAD and the vitamins are coenzymes and electron carriers for diverse metabolic processes including ATP generation, and as mentioned above for converting succinate to fumarate.

Other major changes in the mutants included elevated levels of urea and ornithine. These rose 4.7 and 2.7-fold, respectively in the mtP&E strains, and lesser but statistically significant amounts in the single mtENO and mtPGK knockouts (Fig. 5). An increase in these compounds normally signals increased amino acid catabolism, often to regenerate TCA intermediates (21). It follows that the concentrations of seven amino acids fell in the double knockouts, which are represented by leucine and serine in Fig. 5. The drop in serine could also be attributed to reduced 3-phosphoglycerate, which is the substrate for the PSB pathway. On the other hand, several amino acids rose in the knockouts, such as tyrosine and phenylalanine. An alternative explanation for the increase in ornithine is a bottleneck in its metabolism, since its conversion to other compounds involves an enzyme dependent on pyridoxine, which declined by 80% in the double knockouts (22).

### A loss of PSAT increases serine levels

The PSAT knockouts also resulted in metabolic changes, albeit fewer than observed with the glycolytic gene mutants. Although our initial hypothesis was that eliminating PSAT would cause serine to drop, the opposite was true: serine rose by an average of 89%. Also increasing were α-ketoglutarate by 21% and 3-phosphoserine by 36%. These changes are consistent with the use of the PSB pathway to catabolize serine to generate 3-phosphoglycerate for glycolysis, rather than for making serine. The increases in α-ketoglutarate and 3-phosphoserine were less than that of serine, possibly due to diversions to other pathways such as the TCA cycle for α-ketoglutarate and sulfur assimilation using phosphoserine sulfhydrolase (PITG_06265) for 3-phosphoserine. There was also a modest (23%) but significant increase in thiamine, possibly due to the slower growth of the PSAT knockouts; if growth slows, cells may continue synthesizing vitamins while consuming them more slowly.

Since the media employed for the experiment lacked serine and glycine, the data also support a model in which serine is made primarily from glycine by SHMT (Fig. 1A). Glycine may be made from other amino acids using three pathways in *P. infestans.* These involve glyoxylate transaminase (PITG_11474), threonine dehydrogenase (PITG_05140, PITG_03685), or threonine aldolase (PITG_02857).

### Novel mitochondrial enzymes are not limited to the glycolysis or serine pathways

To assess if additional *P. infestans* enzymes differed from their non-stramenopile counterparts by localizing to mitochondria, all 1524 of its metabolic enzymes were checked for mitochondrial targeting with DeepLoc 2.1, using a conservative cut-off score of 0.875. Of the 224 proteins predicted to be mitochondrial, comparisons were made with proteins from human, fungi, slime molds, apicomplexans, plants, diatoms, brown algae, and other oomycetes. This identified 17 enzymes that are mitochondrial in *P. infestans* but not non-stramenopiles, besides the six glycolytic and two PSB enzymes (Table S3).

Five of these *P. infestans* mitochondrial enzymes resembled the glycolytic pay-off enzymes in being cytosolic in animals, fungi, and slime molds and plastidic in plants. However, only ornithine decarboxylase had both cytosolic and mitochondrial forms in oomycetes *(e.g., Phytophthora* and *Pythium).* Unlike the pay-off enzymes only a cytosolic ornithine decarboxylase was detected in other stramenopiles; this was confirmed based on five diatom and two brown algal genomes. The four other enzymes in this category were exclusively mitochondrial in the oomycetes. Two encoded ribose-phosphate pyrophosphokinase, which is the starting point for purine and pyrimidine biosynthesis. Also fitting this pattern (except being absent from humans) were the two subunits of a bacteria-like glutamate synthase. This is the only type of glutamate synthase made by *Phytophthora,* out of the three classes found across other organisms (23).

Two enzymes were predicted to be exclusively cytoplasmic in all non-stramenopiles but mitochondrial in oomycetes. These were glutamine amidotransferase (GATase; PITG_03467) and glucosamine-fructose-6-phosphate aminotransferase (GFAT; PITG_02122), which both transfer the amide nitrogen of glutamine to another molecule in reactions that also make glutamate. It is notable that the other enzyme that forms glutamate, glutamate synthase, is also mitochondrial while the *P. infestans* enzyme that catabolizes that amino acid, glutamine synthetase, is cytosolic. This may prevent futile cycling.

Several enzymes that were mitochondrial in *P. infestans* and other stramenopiles had a limited distribution in eukaryotes, being found only in plants. These included L-aspartate oxidase (PITG_16480) and quinolinate synthetase (PITG_06783), which comprise a pathway that produces quinolinic acid, which is the precursor for NAD. These are plastidic in plants but absent from most other eukaryotes which instead make quinolinate from tryptophan (24). Other enzymes also lacked orthologs in most eukaryotes but within stramenopiles were only in oomycetes. Two of these, acetate kinase and phosphate acetyltransferase, have been suggested as candidates for HGT from bacteria but were not reported to be mitochondrial (25). These form a pathway for producing acetyl-CoA from acetate or ATP from excess acetyl-CoA. They represent another case beyond the pay-off and PSB systems where oomycetes diversified their metabolism by acquiring multi-enzyme mitochondrial pathways by HGT.

## DISCUSSION

This study has demonstrated that the pay-off and PSB enzymes with the atypical mitochondrial location are required for fitness in *P. infestans*. Single-gene biallelic knockouts of mtPGK and mtENO had limited impact, likely due to flexibility afforded by having dual glycolytic pay-off phases combined with the ability to obtain glycolytic intermediates from other pathways, such as PSB. However, these factors could not overcome the deletion of both mtPGK and mtENO, which severely impaired growth, sporulation, and virulence. Effects of the double deletion were reflected in metabolic imbalances that extended beyond glycolysis to the TCA cycle, urea cycle, amino acids, cofactors, and the adenylate energy currency. In contrast, the influence of the PSAT deletion was more limited although serine levels and growth were affected.

When interpreting the metabolic consequences of the deletions, it is important to consider that our results on synthetic media may not match what occurs *in planta.* Unfortunately, current technologies do not allow pathogen and host metabolites to be measured separately during infection. Nevertheless, we used a medium that was rich in amino acids as these are abundant in plants and are a preferred nitrogen source, and a carbon source, for *P. infestans* (26–28). We omitted serine and glycine to help test the importance of the PSB pathway, but this may have been unnecessary since it appeared to mostly generate 3-phosphoglycerate to fill the pay-off pathway and not to make serine (or ⍺-ketoglutarate). However, in some environments the pathway might be used to create serine, which is an important source of one-carbon units for metabolism (29). This could be addressed by measuring flux using a ^13^C tracer, but the value of performing this in synthetic media is limited. Differences between synthetic media and plants may also explain why the PSAT knockouts reduced growth less in tomato leaves than media. The ability of *P. infestans* to rely on host metabolites may also explain why the defects caused by the PSAT knockouts were less severe than those observed in plants and animals, where deficiencies in that enzyme stunt growth or are fatal (30, 31).

Similar to our work with *P. infestans,* several studies have mutated the plastidic glycolytic enzymes of plants. We are unaware of experiments using double mutants, however deficits in single enzymes had diverse effects ranging from mild phenotypes with enolase to seedling lethality with phosphoglycerate kinase (32, 33). The mildness of many of the effects has been attributed to the exchange of glycolytic intermediates between chloroplast and cytosol by a triose phosphate translocator (33, 34). Comparing our results with *P. infestans* with the plant studies is interesting since the pay-off enzymes occur in two locations in both systems. However, while we propose that the parallel pathways have similar functions in oomycetes, in plants they play distinct roles as the cytosolic pathway is involved in glycolysis while the chloroplast enzymes function in the Calvin cycle.

Our prior phylogenetic analyses were consistent with the acquisition by stramenopiles of most of the mitochondrial pay-off enzymes by endosymbiotic or horizontal gene transfer (E/HGT) from chloroplasts or their ancestor, cyanobacteria (15). Both potential donors have the Calvin cycle (35), but in stramenopiles the enzymes appear to have adapted to a glycolytic role. Since plastid and mitochondrial transfer peptides are similar, it is not surprising that a plastidic protein could adapt to mitochondrial import (36). The PSAT sequence was likely acquired by the same E/HGT event since it and mtPGK are adjacent in the *P. infestans* genome. This apparently also generated the PGDH-PSAT fusion since while its PGDH domain clusters with cytosolic animal proteins, its PSAT domain groups with plastidic orthologs (15). PGDH is N-terminal to the PSAT domain, so the mitochondrial transfer peptide either arose later or evolved from a plastid import sequence that shifted to the N-terminus.

Why the novel pay-off and PSB genes were retained in the stramenopile ancestor is an intriguing question since similar activities were already encoded by its genome. This is distinct from situations where HGT conferred a new function such as the acquisition of anaerobic enzymes by the protist *Giardia* (37) or enzymes with new substrate specificities by the red alga *Galdieria* (38). When an HGT gene is redundant with a native gene, a typical outcome is that one copy becomes specialized or is lost (39). The latter, xenologous gene replacement, applies to the PSB pathway since the cytosolic pathway acquired by descent was lost. With the glycolytic enzymes, dual pay-off phases may have been retained if this increased metabolic output by circumventing the need to import pyruvate into mitochondria, or by doubling the amount of enzyme for the pathways. The latter would resemble the situation in some bacteria where two forms of lysine-tRNA synthetases coexist, one acquired by HGT (40). This might help a plant pathogen like *P. infestans* which needs to adapt to host-imposed stresses and nutrient variability. However, a similar benefit would need to apply to its stramenopile ancestor, which was likely non-pathogenic (41).

Interestingly, acquisition of the mitochondrial pay-off enzymes in the stramenopile ancestor may have driven other evolutionary innovations. These species were recently reported to have evolved proteins capable of transporting triose phosphates from the cytosol across the mitochondrial inner membrane, ostensibly to allow cross-talk between the two pay-off stages (42). Besides highlighting the diversity of eukaryotic metabolism, the novel glycolytic and PSB enzymes and the transporters represent potential targets for inhibitors of oomycetes and other stramenopiles that infect animals (43, 44) or plants (45).

## MATERIALS AND METHODS

### Culturing and life stages of *P. infestans*

Strain 1306 was cultivated at 18°C on rye-sucrose media containing 1.5% agar (46), rye-sucrose broth clarified by centrifugation, or synthetic media adapted from Severino et al. (47). Per liter, this contained 10 g glucose, 5 g sucrose, 1 mg thiamine, 0.4 g KH_2_PO_4_, 0.4 g NaNO_3_, 0.1 g CaCl_2_, 0.1 g MgCO_3_, 0.1 g (NH_4_)_2_SO_4_, 20 mg FeSO_4_·7H₂O, 0.2 g succinate, arginine, glycine, and aspartate, 0.4 g glutamate, 0.15 g mg cysteine, and 0.1 mg of each of the remaining 15 amino acids, pH 5.0. Some experiments omitted glycine and serine as described in the text. Dry weight assays were carried out by inoculating 20 ml of broth with 10^5^ sporangia or, for double mutants, by spreading sporangia on agar overlaid with a 0.4 µm pore polycarbonate membrane. After a brief wash, the mycelia were collected, microwaved to dryness, and weighed. For measuring sporulation, sporangia were released from 100-mm culture plates eight days after inoculation by flooding with 10 ml of water and rubbing with a bent rod. The spores were then counted using a hemocytometer. Zoosporogenesis was induced by incubating sporangia in deionized water for 2.5 hr at 10°C.

Infection assays used leaves from potato cultivar Russet Burbank or tomato cultivar New Yorker. For most assays the leaves were placed on 1.0% water agar, inoculated with two 15 µl droplets of 10^5^/ml sporangia on the midline of their top surfaces, and incubated at 18°C with a 12 hr day-night cycle. Studies of the latent period were initiated by dipping leaves in a solution of zoospores (2×10^4^/ml). Leaves were monitored daily for sporulation. For AUDPC calculations, sporulation on >50% of the leaf surface was scored as 1.0 while sparser sporulation received 0.5. No necrosis was observed since strain 1306 is very biotrophic.

### RNA analysis and bioinformatics

Whether metabolic enzymes from *P. infestans* were mitochondrial was predicted using DeepLoc2.1 (48). Orthologs were extracted from the eggNOG database (49) and filtered to ensure that the proteins had the same specific functional prediction in the Conserved Domain Database (50). Then, their subcellular localizations were assessed using DeepLoc2.1. If that program yielded similar prediction scores for mitochondria and plastid, LOCALIZER served as a tiebreaker (51). When sequences appeared to have truncated N-termini, an effort was made to extend the gene model. Conclusions were also drawn using orthologs from other members of the genus or species group and by phylogenetic analysis.

RNA-seq analysis employed sequence reads that we had prepared as part of other projects (52, 53). Expression calls were made using SystemPipeR with TMM normalization (54). For Fig. 1B, FPKM values were used to allow gene size-corrected comparisons of the mitochondrial and cytosolic forms.

### CRISPR-Cas12a knockouts

Editing was performed as described (55) using pSTUC-1 with a post-transformation incubation temperature of 25°C. Each gene was targeted by three to four gRNAs within or upstream of the catalytic domain (Table S4). The editing plasmid, including Cas12a and a gRNA array, was introduced into *P. infestans* by protoplast transformation. To detect editing, genomic DNA from transformants was analyzed by PCR using the primers in Table S5 and sequenced. Transformants of interest were passed through a single zoospore stage to ensure that they were homokaryons.

### Phosphoglycerate kinase assay

This employed a kit from Abcam (cat. ab252890). In brief, mycelia from 6-day cultures were homogenized in cold assay buffer and debris removed by centrifugation. Reactions were initiated by adding assay buffer (pH 7.2), PGK developer, ATP, NADH, and 3-phosphoglycerate to the extracts (omitted in background wells). Absorbance at 340 nm was recorded every 30 sec for 20 min at 25°C and the change in the last 15 min converted to nmol NADH using a standard curve. Protein concentrations were measured with a dye-binding assay (56) to calculate specific activity.

### Metabolomics

Samples were grown by spreading sporangia on a polycarbonate membrane overlaid on the amino acid-based media lacking glycine and serine, using four replicates per strain. After 5 days at 18°C, the membrane containing the mycelium was removed and washed for 15 sec on a Buchner funnel under mild vacuum. The mycelium was then placed in a cryotube, flash-frozen, lyophilized, ground, and about 10 mg placed in a preweighed vial. After weighing, the powder was extracted with 100 µl per mg of 30:30:20:20 acetonitrile:methanol:isopropanol:water using sonication on ice for 5 min and 90 min of vortexing at 4°C, and clarified by centrifugation. Polar primary metabolites were then quantified by LC-MS by the UC-Riverside Metabolomics Core Facility.

## Acknowledgments

We thank Amancio Jose de Souza and Haiyan Ke for performing the metabolomics analysis. The project was supported by awards to HSJ from the National Science Foundtion.

## Data availability statement

The data that support the findings of this study are in the supplementary files or are available in the NCBI Sequence Read Archive under Bioprojects PRJNA361417 and PRJNA407960.

## Supplementary tables

Table S1. FPKM values for genes shown in Fig. 1B.

Table S2. Metabolomic data for unedited controls and knockout strains.

Table S3. Additional *P. infestans* enzymes with an atypical mitochondrial location.

Table S4. Guide RNAs used for editing.

Table S5. Oligonucleotides used for cloning and PCR.

