## Supplemental Table 3 for "Glycolytic and serine biosynthetic pathways with a novel mitochondrial location contribute to fitness in the oomycete *Phytophthora*"

**Table S3.** Additional P. *infestans* enzymes with atypical subcellular location in mitochondria compared to other eukaryotes

| Gene | Description | DeepLoc score for mitochondria (*P. infestans*) | *Species with cytosolic (C), plastidic (P), or mitochondrial (M) forms* | | | | | | | | | |
|---|---|---|---|---|---|---|---|---|---|---|---|---|
| | | | *Homo sapians* | *Fusarium oxysporum* | *Dictyostelium discoideum* | *Arabidopsis thaliana* | *Chlamydomonas reinhardtii* | *Plasmodium falciparum* | *Ectocarpus siliculosus* | *Phaeodactylum tricornutum* | *Pythium irregulare* | *Phytophthora parasitica* |
| PITG_02122 | glucosamine-fructose-6-phosphate aminotransferase | 0.9259 | C | C | C | C | C | C | P | C | M | M |
| PITG_02303 | ornithine decarboxylase | 0.9203 | C | C | C | P | P, C[2] | C | C | C | M, C[2] | M, C[2] |
| PITG_03467 | glutamine amidotransferase | 0.8817 | nd[1] | C | C | C | C | C | nd | C | M | M |
| PITG_05551 | ribose-phosphate pyrophosphokinase | 0.9508 | C | C | C | P | P | M | P | nd | M | M |
| PITG_05953[5] | acetyldiaminobutyrate dehydratase | 0.9464 | nd | nd | nd | nd | nd | nd | nd | nd | M | M |
| PITG_06783 | quinolinate synthetase A protein | 0.9369 | nd | nd | nd | P | P | nd | M | M | M | M |
| PITG_07380 | glutamate synthase $\alpha$ chain | 0.9192 | nd | C | nd | P | P | C | M | M | M | M |
| PITG_09456[4] | nitroreductase family | 0.9515 | nd | nd | nd | nd | nd | nd | M | M | M | M |
| PITG_09817 | ribose-phosphate pyrophosphokinase | 0.9754 | C | C | C | P | P | M | P | nd | M | M |
| PITG_10301[4] | homoserine O-acetyltransferase | 0.9006 | nd | C[6] | nd | nd | nd | nd | nd | M | M | M |
| PITG_12037 | glutamate synthase $\beta$ chain | 0.9062 | nd | C | C | P | P | C | M | M | M | M |
| PITG_12727 | cysteine synthase | 0.9576 | C | C | C | P, M, C[3] | P, C[3] | nd | C | P, M, C[3] | M | M |
| PITG_13448 | formate dehydrogenase | 0.8726 | nd | C | nd | M | nd | nd | nd | nd | nd | M |
| PITG_15242 | phosphate acetyltransferase | 0.9592 | nd | nd | nd | nd | P | nd | nd | nd | M | M |
| PITG_15243 | phosphate acetyltransferase | 0.9461 | nd | nd | nd | nd | P | nd | nd | nd | M | M |
| PITG_16480 | L-aspartate oxidase | 0.9835 | nd | nd | nd | P | P | nd | M | M | M | M |
| PITG_18248[7] | acetate kinase | 0.9578 | nd | C | nd | nd | C | nd | nd | nd | M | M |

Notes

1. None detected in the indicated species and other members of the genus or group.
2. In addition to the three mitochondrial form, three cytosolic forms were detected in *Chlamydomonas, Phytophthora,* and *Pythium*
3. *Arabidopsis* has two, three, and two plastidic, mitochondrial, and cytosolic forms. *Chlamydomonas* has two plastidic and one cytosolic form. *Phaeodactylum* has one cytosolic and one mitochondrial form.
4. Candidate for horizonal gene transfer from bacteria into stramenopile ancestor.
5. Candidate for horizonal gene transfer from bacteria into oomycete ancestor.
6. Multiple HGT events seem likely during the eukaryotic radiation. In phylogenetic analyses, fungal, oomycete, and diatom clades form three well-separated and well-supported clades with diatoms clustering with one group of bacteria, fungi with a different group, and oomycetes with lower plants *(Chlamydomonas, Emiliania, Physcomitrella, Seliginella).* An ortholog is not found in higher plants.
