## Supplemental Tables 4 and 5 for "Glycolytic and serine biosynthetic pathways with a novel mitochondrial location contribute to fitness in the oomycete *Phytophthora*"

**Table S4.** Guide RNAs (gRNAs) used in this study.

| Gene | Strand | 5’-PAM | gRNA (5’ to 3’) |
| --- | --- | --- | --- |
| PITG_00132 (mtPGK) | +  +  - | TTTA TTTG TTTG | AGAAGAATGTCACGCCCGCCACT GGCGAACTTCTCAGCAGCCCTGT ATGAAGTTAGTCACGCCCTCTGT |
| PITG_14195 (mtENO) | -  +  -  - | TTTG TTTC TTTC TTTG | TGCACGCACAATTCATAAAATGC ACTCTGCGTTTTGCTCCAAGTTC TTCTTGATGACGCTCTTGAGCTG GCGTCGCTGCCGAGAGCCTCTTC |
| PITG_00132 (PGDH-PSAT) | -  -  + | TTTG TTTG TTTG | CCGCTCTCAATGGCTTCCAGCAG TGAACGTCACCTCCTCCTCCAGC AGAAGGAGAACGGCACGTCCAAG |

**Table S5.** Oligonucleotides used in this study.

| Name | Description | Sequence (5’ to 3’) |
| --- | --- | --- |
| PITG_00132_ CRISPRFor, PITG_00132_ CRISPRRev | Annealed to make CRISPR array against mtPGK | AAAATAATTTCTACTAAGTGTAGATAGAAGAATGTCACGCCCGCCACTTAATTTCTACTAAGTGTAGATGGCGAACTTCTCAGCAGCCCTGTTAATTTCTACTAAGTGTAGATATGAAGTTAGTCACGCCCTCTGT,  ATTAACAGAGGGCGTGACTAACTTCATATCTACACTTAGTAGAAATTAACAGGGCTGCTGAGAAGTTCGCCATCTACACTTAGTAGAAATTAAGTGGCGGGCGTGACATTCTTCTATCTACACTTAGTAGAAATTA |
| PITG_14195_ CRISPRFor, PITG_14195_ CRISPRRev | Annealed to make CRISPR array against mtENO | AAAATAATTTCTACTAAGTGTAGATACTCTGCGTTTTGCTCCAAGTTCTAATTTCTACTAAGTGTAGATTGCACGCACAATTCATAAAATGCTAATTTCTACTAAGTGTAGATTTCTTGATGACGCTCTTGAGCTGTAATTTCTACTAAGTGTAGATGCGTCGCTGCCGAGAGCCTCTTC, ATTAGAAGAGGCTCTCGGCAGCGACGCATCTACACTTAGTAGAAATTACAGCTCAAGAGCGTCATCAAGAAATCTACACTTAGTAGAAATTAGCATTTTATGAATTGTGCGTGCAATCTACACTTAGTAGAAATTAGAACTTGGAGCAAAACGCAGAGTATCTACACTTAGTAGAAATTA |
| PITG_00133_ CRISPRFor, PITG_00133_ CRISPRRev | Annealed to make CRISPR array against PGDH-PSAT | AAAATAATTTCTACTAAGTGTAGATCCGCTCTCAATGGCTTCCAGCAGTAATTTCTACTAAGTGTAGATTGAACGTCACCTCCTCCTCCAGCTAATTTCTACTAAGTGTAGATAGAAGGAGAACGGCACGTCCAAG, ATTACTTGGACGTGCCGTTCTCCTTCTATCTACACTTAGTAGAAATTAGCTGGAGGAGGAGGTGACGTTCAATCTACACTTAGTAGAAATTACTGCTGGAAGCCATTGAGAGCGGATCTACACTTAGTAGAAATTA |
| PITG_00132_PCR_F,  PITG_00132_PCR_R | PCR to detect editing of mtPGK | TCTCGCGTATGAGCATGCACTTG, ATGATCTTGTCGCACTTTTGTAG |
| PITG_14195_PCR_F,  PITG_14195_PCR_R | PCR to detect editing of mtENO | CAACATGGCGATTTTCTGAAAC, AGCCCGTTTCAATACAGTTATT |
| PITG_00133_PCR_F,  PITG_00133_PCR_R | PCR to detect editing of PGDH-PSAT | CTGCAACGCCTTCGGAATGAAT, TCACGCGCCAAGCCGAAGTTAC |
